# Mycobacterial Metabolic Model Development for Drug Target Identification

**DOI:** 10.1101/2023.03.31.534705

**Authors:** Bridget P. Bannerman, Alex Oarga, Jorge Júlvez

**Affiliations:** University of Cambridge, Cambridge, UK; Department of Computer Science and Systems Engineering, University of Zaragoza, Zaragoza, Spain; Science Resources Foundation, UK

**Keywords:** *M. abscessus*, *M. leprae*, *M. tuberculosis*, drug targets, bottleneck reactions

## Abstract

Antibiotic resistance is increasing at an alarming rate, and three related mycobacteria are the source of widespread infections in humans. According to the World Health Organization, *Mycobacterium leprae*, which causes leprosy, is still endemic in tropical countries; Mycobacterium tuberculosis is the second largest infectious disease killer worldwide after COVID-19; *Mycobacteroides abscessus*, a group of non-tuberculous mycobacteria, causes lung infections and other health-care-associated infections in humans. Due to the rise in resistance to common antibacterial drugs, it is critical to develop alternatives to traditional treatment procedures. Furthermore, an understanding of the biochemical mechanisms underlying pathogenic evolution is important for the treatment and management of these diseased conditions.

In this study, metabolic models have been developed for two bacterial pathogens, *M. leprae*, and *M. abscessus*, and a new computational tool has been used to identify potential drug targets, which are referred to as bottleneck reactions. The genes, reactions, and pathways in each of these organisms have been highlighted; the potential drug targets can be further explored as broad-spectrum antibacterials and the unique drug targets to each pathogen are significant for precision medicine initiatives.

The models and associating datasets are available in GigaScience and the following repositories:

- *M. abscessus*
  - Biomodels: https://www.ebi.ac.uk/biomodels/MODEL2203300002
  - https://www.patmedb.org/Bacteria/Mabscessus
- *M. leprae*
  - Biomodels: https://www.ebi.ac.uk/biomodels/MODEL2203300001
  - https://www.patmedb.org/Bacteria/Mleprae

## DATA DESCRIPTION

### Context

*M. leprae* and *M. tuberculosis* are two related pathogenic mycobacteria responsible for leprosy and tuberculosis in humans. Another related mycobacterium, *M. abscessus*, causes opportunistic infections in healthcare-related settings.

The previous analyses of the metabolic models of *M. tuberculosis* have supported studies demonstrating the evolutionary drivers of antibiotic resistance and the identification of novel drug targets against mycobacteria [Kavvas *et al*., 2018, Bannerman *et al*., 2019].

Here we demonstrate the newly built genome-scale metabolic models of *M. leprae* and *M. abscessus*, including the curation, simulation, and model optimization strategies [Ebrahim *et al*., 2013, Oarga *et al*., 2020, Bannerman *et al*., 2021]. To ensure the development of standardised metabolic models for the global systems biology community, we have implemented the recently released community standards and used the MEMOTE quality control software to evaluate our models [Carey *et al*., 2020, Lieven *et al*., 2020].

## METHODS

### GEM reconstruction, curation, and simulation

Automated draft reconstructions of *M. leprae* and *M. abscessus* were downloaded from BioModels and evaluated them against other organism-specific databases, such as BioCyc and KEGG [Caspi, 2014; Kanehisa, 2021]. COBRApy, a Python toolbox for the construction, manipulation, and analysis of constraint-based models [Ebrahim *et al*., 2013] and GNU Linear Programming Kit (GLPK), a software package which solves efficiently large-scale linear programming problems (https://www.gnu.org/software/glpk/) were used to manipulate and simulate the models.

To create the new genome-scale metabolic models (GEM) of *M. abscessus* and *M. leprae*, additional reactions, gene-to-reaction associations, and pathways (that were not in the automated model) were integrated from Kegg and BioCyc [Caspi, 2014; Kanehisa, 2021]. The annotation of genes and metabolites were improved by comparing and transferring annotations from the related *M. tuberculosis* models in iEKVIII model and BioCyc database [Caspi, 2014; Kavvas *et al*., 2018]. Further improvements to the models were made by comparing them with the compound formula and charge from the Metanetx database and by mapping the genes and reactions of the GEMS to the BiGG, ChEBI, KEGG, and MetaCyc, databases [Hastings *et al*., 2016; Norsigian *et al*., 2020; Moretti *et al*, 2021]. Figure 1 describes the full process of the mycobacterial metabolic models (*i*Mab22 and *i*Mlep22) of the pathogens, *M. abscessus* and *M. leprae*. The revised model reconstructions of *M. leprae* (*i*Mlep22) and *M. abscessus* (*i*Mab22) can be instantiated without error on the COBRA software (version 0.16.0) [Ebrahim *et al*, 2013].

**Figure 1:**
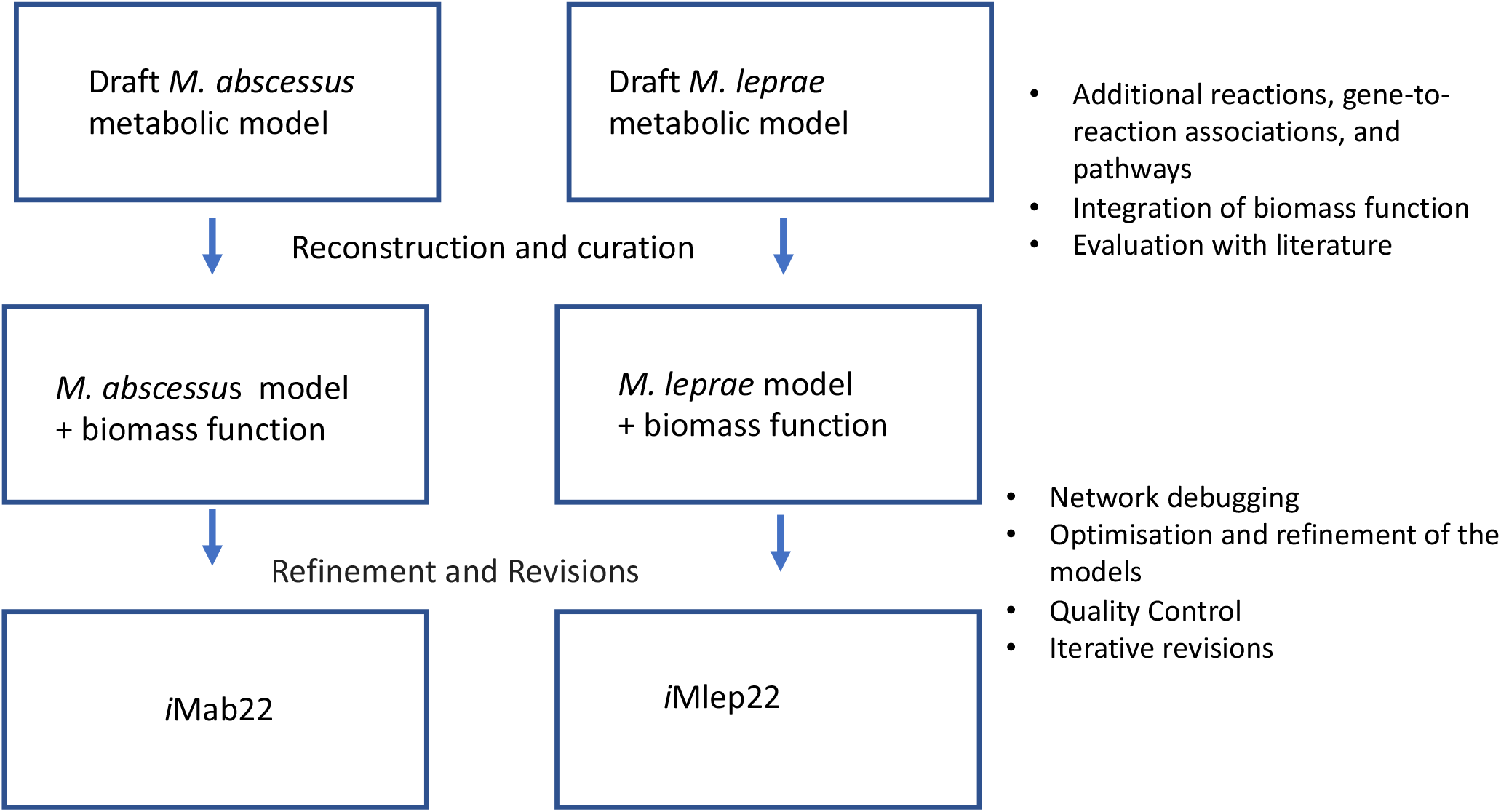
Mycobacterial Metabolic Model Development for Drug Target Identification: *M. abscessus* and *M. leprae*

### BIOMASS reactions for *M. leprae* and *M. abscessus*

We generated biomass reactions for *M. abscessu*s and *M. leprae* using the methodology in the pathway tools software [Karp *et al*., 2019] and the corresponding BioCyc databases [Karp *et al*., 2019]. The software tool, *findCPcli*, which implements the computational method for identifying bottleneck reactions as drug targets are available at https://github.com/findCP/findCPcli.

## DATA VALIDATION AND QUALITY CONTROL

### Model optimisation

The dead-end metabolite reactions that were previously present in the automated model were eliminated to enhance the model’s quality. The model was then iteratively evaluated, considering model-specific reactions for the *M. abscessus* and *M. leprae* organisms and comparisons to the BioCyc, Kegg, MetaNetX 4.2, BiGG, and ChEBI databases [Caspi *et al*., 2014; Hastings *et al*., 2016; Norsigian *et al*., 2020; Moretti *et al*., 2021].

To improve the quality of the model, the reactions with dead-end metabolites, previously found in the automated models were removed. MEMOTE, a standardised genome-scale metabolic model testing programme was used to undertake quality control checks during the model’s iterations and optimization [Leiven *et al*., 2020]. In the process, SBO annotations and gene annotations from the KEGG database and 728 new formulae from the MetaNetX database were added [Moretti *et al*., 2021]. As a result, the Memote score increased from 49% to 63% on the *M. leprae* model and from 48% to 66% on the *M. abscessus* model.

The new GEMS for *M. abscessus* and *M. leprae* are encoded in the Systems Biology Markup Language (SBML) [Keating *et al*., 2020] and designated as follows:

*i*Mlep22 for the *M. leprae* model: i for in silico, Mlep for *M. leprae*, and published in 2022. *i*Mlep22 consists of 5625 reactions, 4016 metabolites, 871 genes.

*i*Mab22 for the *M. abscessus* model: i for in silico, Mab for *M. abscessus*, and published in 2022. *i*Mab22 consists of 8580 reactions, 6273 metabolites, and 1837 genes.

Standardisation and curation have been done according to the community standards for the development of metabolic models as described in Carey *et al*., 2020 and Lieven *et al*., 2020 to produce gold standard metabolic network reconstructions of *M. abscessus* and *M. leprae*. The development of *M. abscessus* and *M. leprae* (*i*Mab22 and *i*Mlep22) metabolic models is illustrated in Figure 1, and the overall capability of the models has been summarised in Figure 2.

**Figure 2:**
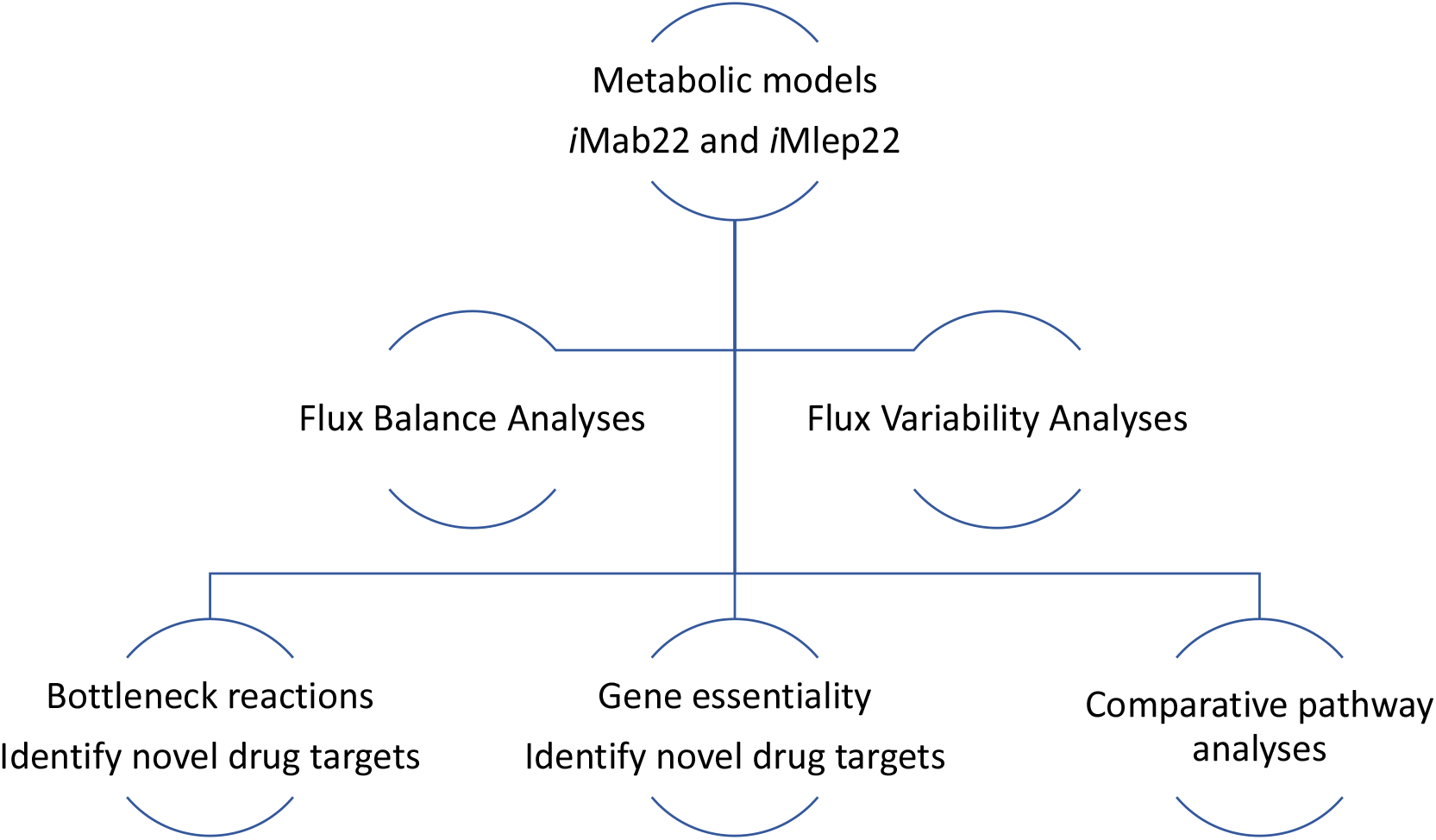
Overall capability of the models : *M. abscessus* (*i*Mab22) and *M. leprae* and (*i*Mlep22)

### Comparative analysis

*M. abscessus*, an opportunistic pathogen, has a large genome size of 5.1 MB (Ripoll *et al*. 2009) with more biochemical and bottleneck reactions in the *i*Mab22 model compared to the obligate pathogens *M. tuberculosis* with a genome size of 4.4MB and *M. leprae*, with an even smaller genome size of 3.3 bp [Singh & Cole, 2011]. The fewer metabolic and bottleneck reactions in the *i*Mlep22 model can be attributed to the reductive evolution that occurred in *M. leprae* and the subsequent loss of genes. An illustration of the distribution of unique enzymes and bottleneck reactions in each of the models (*i*Mab22 and *i*Mlep22) in comparison with each other and with *M. tuberculosis* is demonstrated in Figure 3. Alternative enzymes are not included in this analysis because they catalyse the same biochemical reactions and do not fit into the category of bottleneck reactions, which are defined as unique reactions responsible for the survival and growth of the organisms in the metabolic network [Bannerman *et al*., 2021; Yeh *et al*., 2004; Oarga *et al*., 2020].

**Figure 3:**
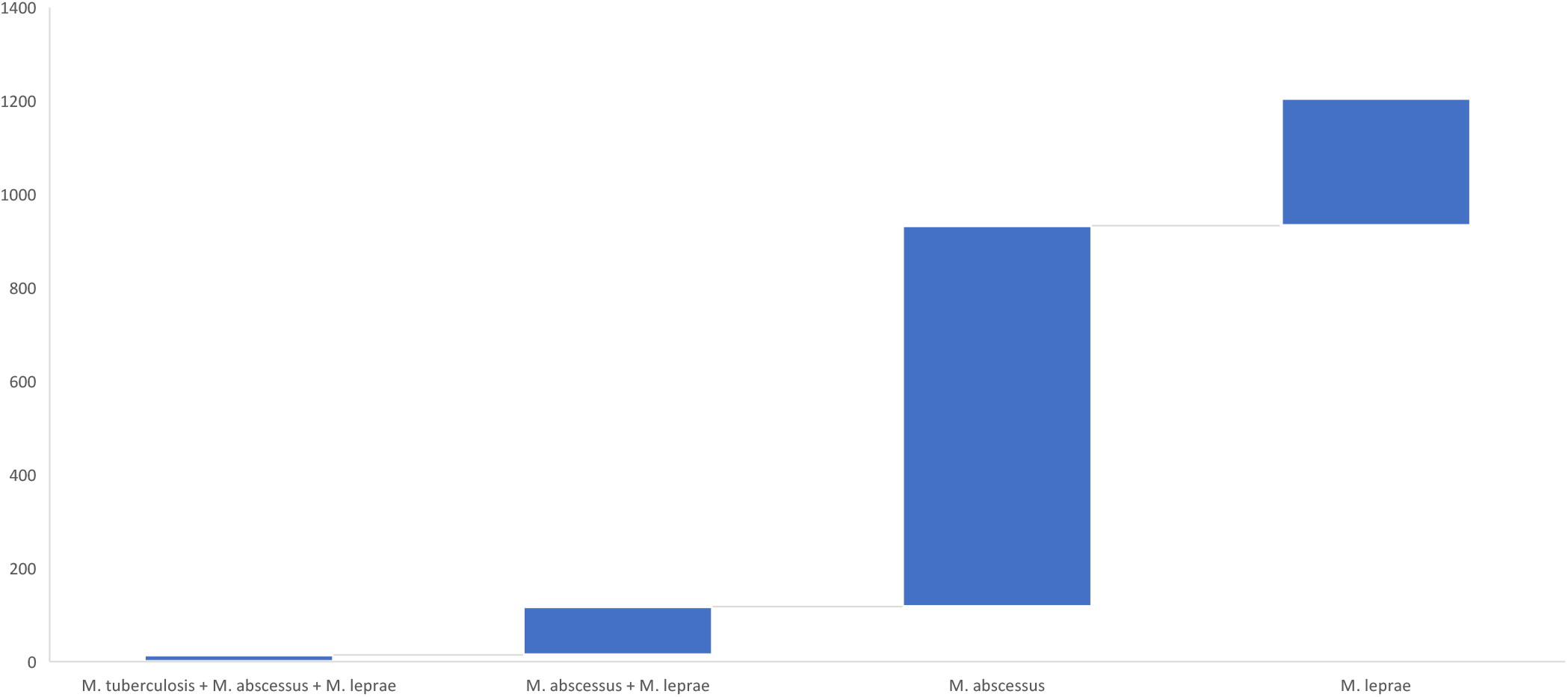
Unique/bottleneck reactions in *M. abscessus, M. leprae* in relation to *M. tuberculosis*. Figure 3b: 1. The number of unique/bottle neck reactions in *M. abscessus* and *M. leprae* 2. The number of bottleneck reactions common to *M. abscessus* and *M. leprae* 3. The number of bottleneck common to *M. abscessus, M. leprae* and *M. tuberculosis*

### Re-use potential

The standardised genomic scale metabolic models for *M. leprae* (*i*Mlep22) and *M. abscessus* (*i*Mab22) have been developed using the systems biology community standards for quality control and evaluation of the models [Carey *et al*., 2020, Lieven *et al*., 2020] and available for reuse by the global scientific community. The models can be retrieved from:

*M. abscessus*

https://www.ebi.ac.uk/biomodels/MODEL2203300002 and

https://www.patmedb.org/Bacteria/Mabscessus.

*M. leprae*

https://www.ebi.ac.uk/biomodels/MODEL2203300001

https://www.patmedb.org/Bacteria/Mleprae

*The findCPcli* tool used for analysing the models can be retrieved from biotools (https://bio.tools/findcpcli) and

SciCruch (https://scicrunch.org/resources/about/registry/SCR_023391).

## Supporting information

Supplemental file: Metabolic model M. abscessus

Supplemental file: Metabolic model M. leprae

## Notes

https://www.ebi.ac.uk/biomodels/MODEL2203300002

https://www.patmedb.org/Bacteria/Mabscessus

https://www.ebi.ac.uk/biomodels/MODEL2203300001

https://www.patmedb.org/Bacteria/Mleprae

https://bio.tools/findcpcli)

https://scicrunch.org/resources/about/registry/SCR_023391

### Competing Interest Statement

The authors have declared no competing interest.

